# Enhanced recovery of CD9-positive extracellular vesicles from human specimens by chelating reagent

**DOI:** 10.1101/2020.06.17.155861

**Authors:** Ayako Kurimoto, Yuki Kawasaki, Toshiki Ueda, Tatsutoshi Inuzuka

## Abstract

Extracellular vesicles (EVs) have gained attention as potential targets of early diagnostics and prognosis in the field of liquid biopsy. Despite clinical potentials, the best method to isolate EVs from specimens remains controversial due to low purity, low specificity, and lack of reproducibility with current isolation methods. Here we show that a chelating reagent enhances the recovery efficiency of EVs from crude biological samples by immunoprecipitation using an anti-CD9 antibody. Proteomic and western blotting analyses show that the EVs isolated using the chelating reagent contain a wider variety of proteins than those isolated with PBS.

## Introduction

EVs are secreted from all types of cells and found in most body fluids, including blood, urine, and saliva(1–3). Several functions of EVs have been reported e.g., intercellular interaction and cancer metastasis; however, the exact biological role of EVs is unclear.

In addition, EVs contain several biologically active molecules such as proteins, mRNA, miRNA, long non-coding RNA (lncRNA), DNA, and phospholipids, which reflect states of cells or organs (4,5). Such biomolecules found in EVs are expected to serve as diagnostic biomarkers or prognostic markers for various diseases in clinical fields.

Despite the high clinical need, EV-based liquid biopsy has not been widely used yet due to a lack of standardized isolation methods from specimens. The most commonly used techniques for EV isolation are ultracentrifugation (UC) and polymer-based precipitation. UC and polymer-based precipitation methods, however, cause aggregation of EVs (6). Furthermore, these techniques are compromised by low selectivity: EVs include microvesicles, apoptotic bodies, and exosomes (7), and such heterogeneity leads to the lack of reproducibility and reliability, hampering clinical utilization. Recent studies suggested that the surface of EVs is positively charged, and it attracts negatively charged molecules such as DNA- and RNA-binding proteins are found on surface of EVs (8).

Rapid and selective isolation of EVs is possible by immunoprecipitation (IP) methods using antibodies against surface marker proteins of interest. Hence, we employed IP method based on surface marker proteins for rapid and specific isolation of EVs. Though, the recovery rate of EVs by IP method is not enough for clinical use, here we report that IP method with a chelating reagent improves yields and purity of EVs from multiple human specimens. The isolated EVs could be used for quantitative and qualitative proteomic analyses with LC-MS/MS. We report that IP method with a chelating reagent improves yields and purity of EVs from multiple human specimens.

## Materials and Methods

### Materials

NIH-H1299 cell lines were obtained from ATCC. Minimum Essential Medium (MEM) Non-essential Amino Acids Solution and Penicillin-Streptomycin solution (x100) were purchased from FUJIFILM Wako Pure Chemical Corporation. Fetal Bovine Serum (FBS) was purchased from Gibco, Thermo Fisher Scientific Inc. Ethylenediaminetetraacetic acid (EDTA) and glycoletherdiamine-N,N,N’,N’-tetraacetic acid (EGTA) were purchased from Dojindo Molecular Technologies, Inc. 1,4-Dithiothreitol (DTT) was purchased from FUJIFILM Wako Pure Chemical Corporation. Anti-CD81 antibody (SHI-EXO-MO3), and pooled human serum were purchased from Cosmo Bio Co., Ltd. A matched set of specimens was purchased from ProMedDx, LLC. MagCapture Exosome Isolation Kit PS was purchased from FUJIFILM Wako Pure Chemical Corporation. Total Exosome Isolation Reagent (from serum) was purchased from Thermo Fisher Scientific Inc. The following antibodies were used for immunostaining: Goat anti-human albumin antibody HRP Conjugated (A80-129P, Bethyl Laboratories, Inc.), goat anti-human IgG H&L (HRP) (ab6858, Abcam plc), anti-apolipoprotein E antibody (Biotin) (ab24274, Abcam plc). Volunteers for saliva and urine samples were recruited from Miraca Research Institute G.K., and the study was conducted with the approval (18-30) by the institutional review board of Miraca Holdings, Inc. Urine and saliva samples were diluted 3-fold with PBS or EDEG.

### Preparation of serum EVs by UC

Serum (1 mL) was centrifuged at 15,000 x g for 15min at 4°C, and the supernatant was ultracentrifuged at 100,000 x g for 1 h at 4°C by himac (Koki Holdings Co., Ltd.) with S55A2 rotor (k-factor: 162). The pellet was resuspended in 1 mL of PBS or EDEG (50 mM EDTA and 50 mM EGTA in PBS), followed by ultracentrifugation at 100,000 x g for 1 h at 4°C to eliminate other contaminating molecules. Resulting EVs were resuspended in PBS and stored at 4°C until use.

### EV isolation from culture media by UC

NIH-H1299 cells were grown in 10% FBS, MEM Non-essential Amino Acids Solution (x100), and Penicillin-Streptomycin solution (x100). Subconfluent cells were washed with PBS twice and grown for 72 h in FBS-free medium before EV isolation. The cell culture supernatants were collected and filtered through 0.22 μm pore membranes using the vacuum filter/storage bottle system (Corning Incorporated) to remove large contaminating vesicles. The obtained conditioned medium was centrifuged at 2,000 x g for 5 min at 4°C, followed by concentration using Amicon Ultra-15 Centrifugal Filter Unit (100 kDa cutoff, Merck KGaA). The concentrated medium (6 mL) was centrifuged at 15,000 x g for 15min at 4°C, and the supernatant was followed by ultracentrifugation at 100,000 x g for 1 h at 4°C by himac (Koki Holdings Co., Ltd.) with an S55A2 rotor (k-factor: 161.6). The pellet was resuspended in 100 μL of PBS or EDEG (50 mM EDTA and 50 mM EGTA in PBS), followed by ultracentrifugation at 100,000 x g for 1 h at 4°C. The resulting pellet that contains EVs was resuspended in PBS and stored at 4°C until use. The protein concentration of the purified EVs was determined by Qubit protein assay kit (Thermo Fisher Scientific Inc.).

### Polymer-based precipitation

Total Exosome Isolation Reagent (from serum) was used according to the manufacturer’s instruction. Frozen pooled serum was thawed at room temperature and centrifuged for 30 min at 2000 x g at 4°C. The supernatant (1 mL) was mixed with 60 μL of Total Exosome Isolation Reagent (from serum). The sample was incubated at 4°C for 30 min. The mixture was centrifuged at 10,000 x g at 4°C and the obtained pellet was resuspended in PBS. The pellet containing EVs was stored at 4°C until use.

### Preparation of antibodies against CD9 and CD63

Antibodies against CD9 and CD63 were prepared as described previously (9) with the modification that 50 μg of EVs from NIH-H1299 obtained by ultracentrifugation method was used as a the source of antigen. Biotinylations of the anti-CD9 monoclonal antibody and anti-CD63 monoclonal antibody were performed using EZ-Link™ Sulfo-NHS-LC-Biotin (Thermo Fisher Scientific Inc.) according to the manufacturer’s protocol.

### IP of EVs

Anti-CD9 antibody coupled with Dynabeads M-280 Tosylactivated (Thermo Fisher Scientific Inc.) was added to matched set specimens (serum and plasma), urine, and saliva that were diluted 1:3 with EDEG or PBS, followed by incubation on a rotator at 4°C for 18 h. The beads were washed three times with PBS and stored at 4°C until further analysis.

### Western blotting analysis

The antibodies were diluted with Can Get Signal (TOYOBO Co., LTD.) to 1 μg/mL. Immunocaptured EVs on Dynabeads M-280 Tosylactivated were lysed by 4x Laemmli Sample Buffer (Bio-Rad Laboratories, Inc.) under nonreducing condition (without DTT) for CD9, CD63, and CD81 or reducing condition (supplemented with 50 mM DTT) for other proteins followed by boiling for 5 min at 96°C. The obtained protein samples were separated by SDS-PAGE and then transferred to PVDF membranes (ATTO Corporation.). After blocking with Blocking One (NACALAI TESQUE, INC.), the membranes were incubated with primary antibodies followed by incubation with secondary antibodies for 1 h at room temperature. The membranes were washed with PBS with Tween® 20 (TAKARA BIO INC.) three times and incubated with secondary antibodies for 1 h at room temperature. After washing, ECL^™^ Select Western Blotting Detection Reagent (GE Healthcare, GENERAL ELECRIC COMPANY) was added to the membrane. Protein was detected by using ImageQuant™ LAS 500 imager (GE Healthcare, GENERAL ELECTRIC COMPANY).

### Preparation of peptide

Pelleted EVs were lysed with 40 μL of 1% *Rapi*Gest™ SF (Waters Corporation) in 50 mM ammonium bicarbonate (Honeywell Fluka™, Thermo Fisher Scientific, Inc.) supplemented with 50 mM DTT, followed by incubation at 60°C for 30 min. After being allowed to cool down to room temperature, the lysed samples were added by 4 μL of 150 mM 2-iodoacetamide and incubated at room temperature for 30 min in the dark. The lysates were incubated with 1 μg/mL of Trypsin/Lys-C Mix, Mass Spec Grade (Promega Corporation) at 37°C overnight. To break down the detergent, 4 μL of 10% of trifluoroacetic acid (Thermo Fisher Scientific, Inc.) was added to the digested mixture and incubated at 37 °C for 30min. After centrifugation at 13,000 x g for 10 min, the supernatant was collected and lyophilized with miVac system (Genevac Ltd) and desalted with Pierce C-18 Spin Columns according to the manufacturer’s instruction. Obtained peptides were eluted with 70% acetonitrile, followed by lyophilization, and stored at -80 °C until use.

### Proteomic analysis with LC-MS

Obtained peptides were reconstituted by 20ul of water containing 0.1% formic acid (FA) (Fisher Chemical, Thermo Fisher Scientific, Inc.). The proteomic analysis of the peptides was carried out on Q Exactive (Thermo Fisher Scientific, Inc.) equipped with UltiMate 3000 Nano LC Systems (Thermo Fisher Scientific, Inc.). Peptide sample (2 μL) was injected onto an Acclaim PepMap 1000 trap column (75 μm × 2 cm, nanoViper C18 3 μm, 100Å, Thermo Fisher Scientific) which was heated to 40 °C in a chamber which was connected to a C18 reverse-phase Aurora UHPLC Emitter Column with nano Zero & Captive Spray Insert (75 μm × 25 cm, Ion Opticks Pty Ltd) using Dreamspray interface (AMR INCORPORATED). Nano pump flow rate was set to 250 nL/min with 170 min gradient, where the mobile phases were A (0.1% FA in water, Fisher Chemical, Thermo Fisher Scientific, Inc.) and B (0.1% FA in acetonitrile, Fisher Chemical, Thermo Fisher Scientific, Inc.). The chromatography gradient was designed to provide a linear increase 0-8 min at 2% B, 8-15 min from 2% B to 15% B, 15-149 min from 15% B to 40% B, 149-150 min from 40% B to 95% B, wash, 8 min and 11 min equilibrium. The data-dependent acquisition was performed in positive ion mode. Mass spectrometer parameters were as follows: MS full scan from *m/z* 350–1500 at a resolution of 70,000, AGC target 3 x e6, maximum injection time 100 ms, and dd-MS2/dd-SIM parameters included AGC target: 1 x e5, maximum injection time: 120 ms, TopN: 10, and isolation window 1.6 *m/z*. dd settings were Minimum AGC of 2.5 x e3, Charge exclusion unassigned, 1, 7, 8, >8, and Dynamic exclusion was set to 30 s.

All MS/MS samples were analyzed using Proteome Discoverer 2.2.0.388, which was set up to search UniProt-human.fasta (downloaded January 2019). Proteins were identified using the following parameters: Precursor mass tolerance: 10ppm, Fragment mass tolerance: 0.02 Da, Max missed cleavage sites: 2, Dynamic modification: Oxidation / + 15.995 Da (M), Dynamic modifications (protein terminus): N-Terminal Modification: Acetyl/ +42.011 Da (N-Terminus), Static Modification: Carbamidomethyl / +57.021 Da (C). Target FDR (Strict): 0.01, Target FDR (Relaxed): 0.05.

### Statistical analysis

For comparison between EVs that were CD9-immunoprecipitated with or without EDEG, proteins were identified in at least three runs and quantified with label-free quantification (LFQ) values using Proteome Discoverer. LFQ parameters were set to Normalization mode: Total peptide amount, Imputation mode: missing value: Low abundance resampling, Ratio calculation: Summed abundance based. Median of obtained normalized abundances of each identified protein, medians of proteins between two sample groups, log2, -log10, p-value, and fold change were calculated using Microsoft Excel and visualized by Volcano plots using R (version 3.5.3).

## Results

### Chelating reagent enhanced the efficiency of IP in serum, plasma, saliva, and urine samples using anti-CD9 antibody

We have compared the yields of immunocapture using the anti-CD9 antibody in matched set specimens. Western blot analysis showed that the CD9 signal of EDTA plasma was the strongest in comparison to serum and plasma containing other anticoagulants e.g., heparin, acid-citrate-dextrose, citrate phosphate dextrose, and sodium citrate (Fig.1 A). Based on the assumption that chelating reagent would enhance the efficiency of immunocapture using the anti-CD9 antibody, we performed immunoprecipitation of CD9 positive vesicles with EDEG reagent (50mM EDTA and 50mM EGTA in PBS). As we have expected, the yields of immunocaptured CD9 with EDEG reagent was improved compared with PBS dilution specimens (Fig.1 B). The yields of CD9 from urine or saliva were slightly improved (Fig.1 C). These differences might reflect the crudeness of specimens, that is, saliva and urine contained fewer contaminant proteins compared to serum and plasma(10).

**Figure 1.**
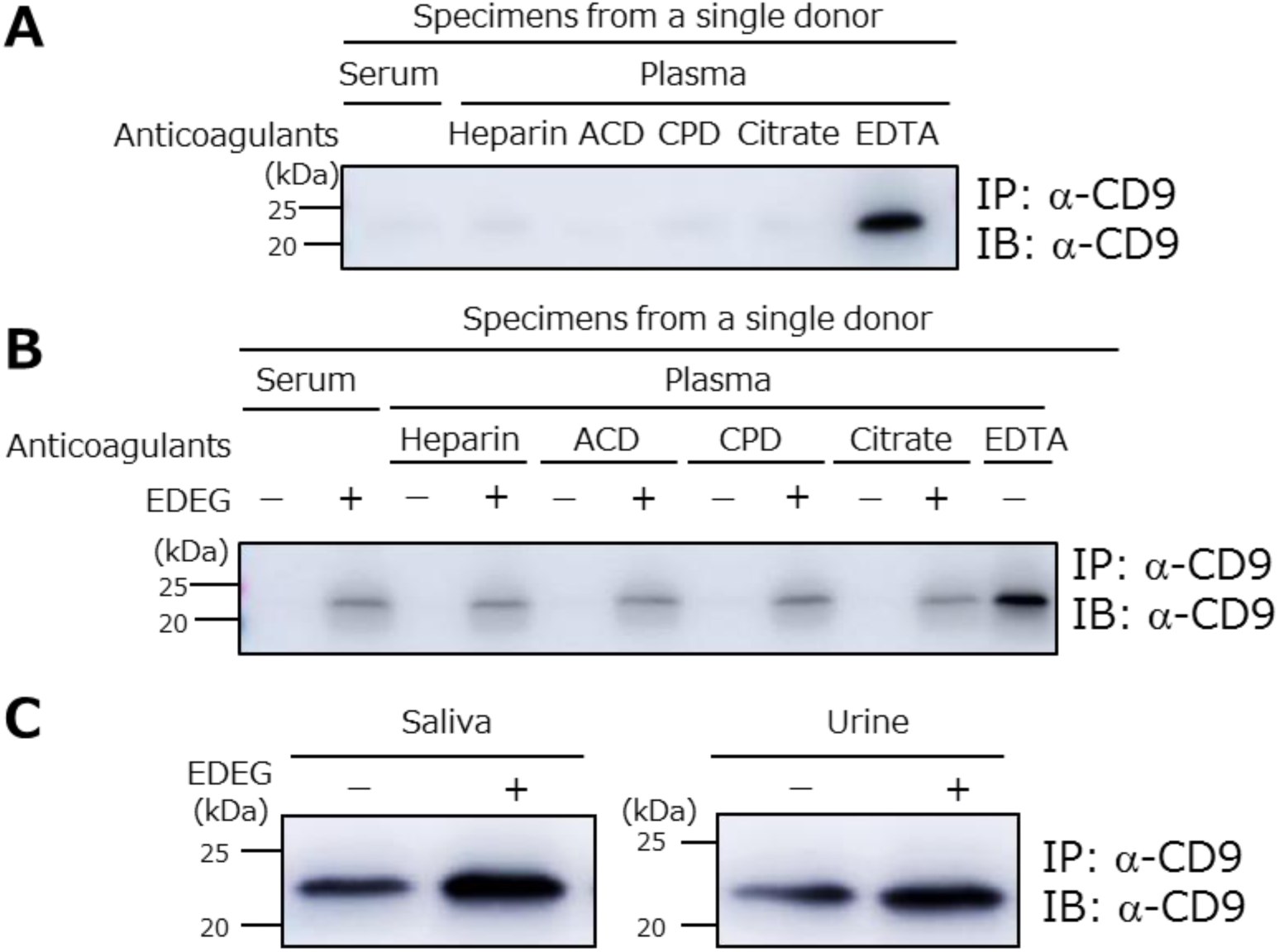
Chelating reagent EDEG enhanced the recovery rate of CD9 positive EVs from serum, saliva, urine, and plasma. (A) Immunoprecipitated CD9 positive EVs from matched set specimens, including serum and plasma containing different anticoagulants, heparin, acid-citrate-dextrose (ACD), citrate phosphate dextrose (CPD), sodium citrate (citrate), and EDTA were detected by anti-CD9 antibody. (B) Western blotting analysis of immunoprecipitated CD9 positive EVs from matched specimen with or without EDEG were performed. (C) Saliva and urine with or without EDEG were subjected to western blot analysis by anti-CD9 antibody. Each specimen used for EV isolation was 300 μL.

### The purity of immunocaptured CD9 positive vesicles was higher using EDEG reagent than immunocapture with PBS, ultracentrifugation method, and polymer-based precipitation

We carried out a series of western blot analyses for serum EVs isolated by Ultracentrifugation (UC), Total Exosome Isolation Reagent (polymer-based precipitation), and immunoprecipitation with an anti-CD9 antibody in EDEG-treated or PBS-treated specimens (EDEG-CD9IP and PBS-CD9IP, respectively). EVs isolated with UC or polymer-based precipitation method were shown to contain both EV-associated proteins (CD9, CD63, and CD81) and non-EV marker proteins (HSA, HIGG, and ApoE) (Fig.2). These results suggest that the EVs isolated using these methods contain a substantial amount of contaminant proteins, which is in line with the previous report (3). On the other hand, EVs in EDEG-CD9IP showed a dramatic reduction in the amount of non-EV marker proteins compared with UC and polymer-based purification. In addition, the amount of EV-associated proteins detected in EDEG-CD9IP was greater than that in PBS-CD9IP. These results indicate that EDEG reagent enables the isolation of highly pure EVs with increased efficiency.

**Figure 2.**
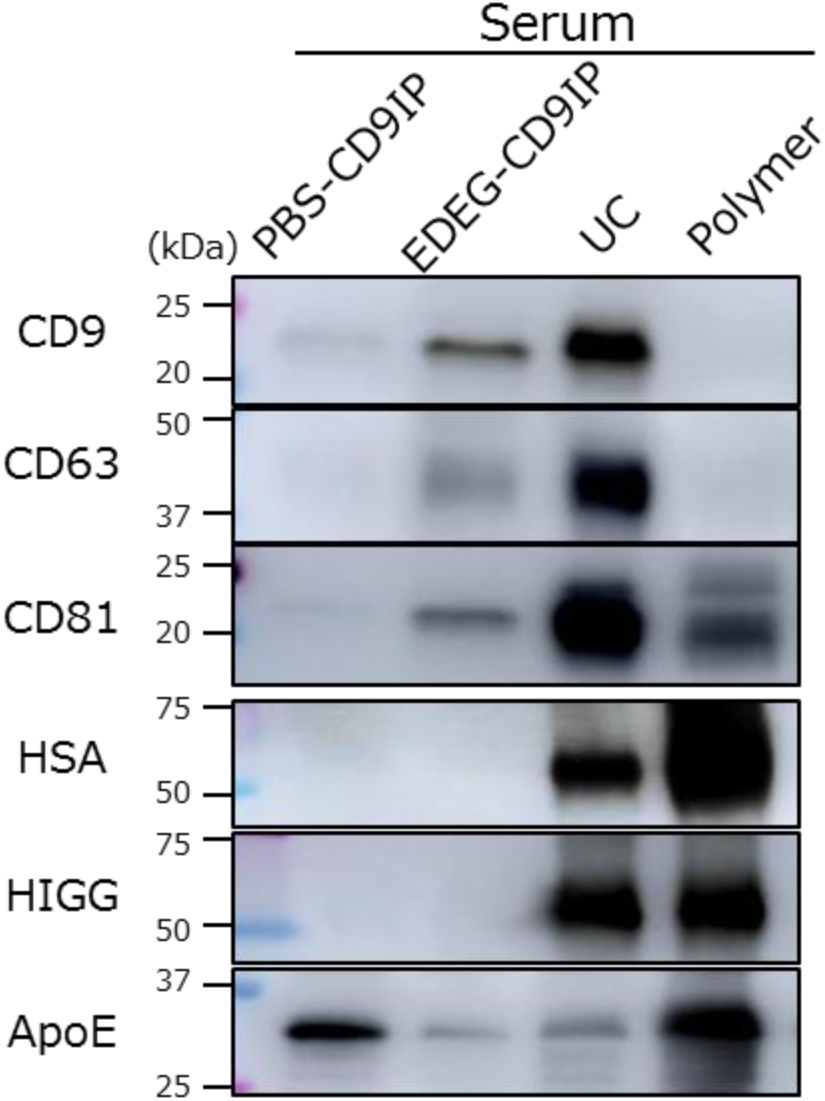
Western blot analysis of serum EVs isolated by CD9 immunoaffinity. PBS-CD9IP, EDEG-CD9IP, UC, and Polymer represent EVs isolated by IP method with CD9 antibody using PBS, using EDEG, ultracentrifugation method, and polymer-based precipitation, respoctively. Abbreviations: HSA, Human serum albumin; HIGG, human immunoglobulin; ApoE, apolipoprotein E.

### Proteomic profiling of CD9 immunoprecipitated EVs from serum diluted with PBS or EDEG reagent

We have carried out proteomic analysis of CD9-immunoprecipitated EVs by LC-MS/MS. Proteomic analysis of EVs with and without EDEG reagent identified 159 proteins and 106 proteins, respectively. Here, EV markers such as CD9 and CD81 were identified only in EDEG diluted serum (Fig.3 A). LFQ analysis was performed to quantify each identified protein. Volcano plot analysis showed EVs immunoprecipitated without EDEG reagent (PBS) contained abundant complement proteins compared to those with EDEG reagent (Fig. 3B). Also, the EVs without EDEG reagent contained complement-related proteins such as properdin and complement C1r, and calcium-binding proteins (i.e. sorcin) (supplementary Table.1). These results suggested that EVs isolated by IP with the EDEG reagent give us access to more potential proteins for clinical diagnosis.

**Figure 3.**
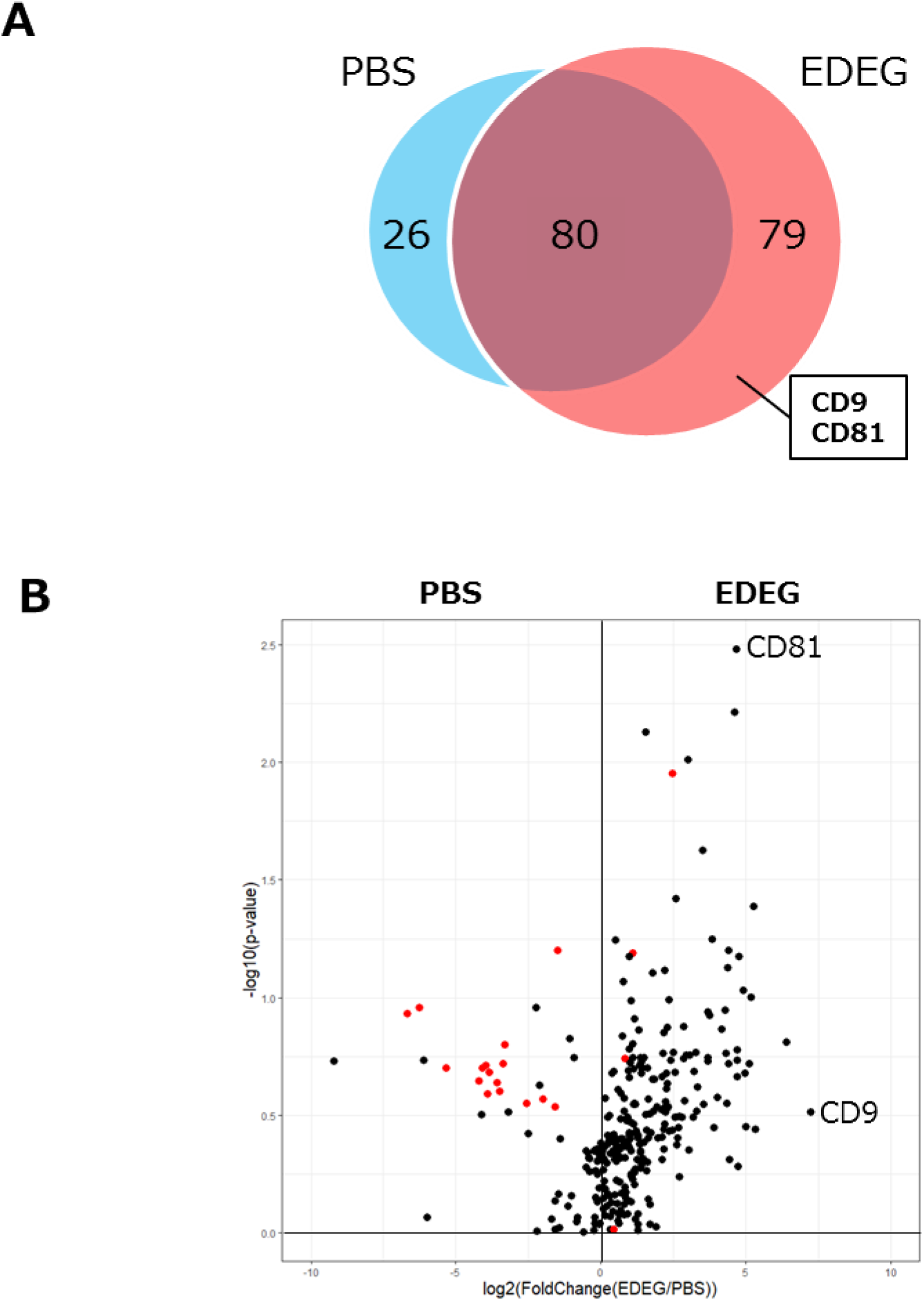
LC-MS analysis of immunocaptured EVs using anti-CD9 antibody from serum diluted with PBS or EDEG reagent. A. Venn diagram displaying proteins immunocaptured CD9 positive EVs from serum diluted with PBS or EDEG. B. Volcano plot analysis. Complement proteins and tetraspanins of interest are highlighted in red.

## Discussion

Although a large number of EV purification methods and kits are now available, it has been demonstrated that non-EV associated proteins and molecules are co-isolated as contaminants (3). In this report, we have shown that IP method using a chelating reagent increases the purity and the yield of CD9 positive vesicles from multiple body fluids, such as serum, plasma, urine, and saliva.

In addition, EVs are heterogeneous vesicles composed of apoptotic bodies, microvesicles, and exosomes(7). The ratio of EV subpopulations and the recovery rate of each subpopulation depends on isolation methods and conditions of biofluids, both of which account for the low repeatability of EV separation.

Accordingly, we have developed a method to improve yields and purity of EVs isolated from multiple specimens for clinical use. In this study, we employed an immunocapture method to isolate EVs with high specificity. As shown in Fig.1A, the amount of recovered CD9 positive EVs varies depending on the added anticoagulants, and chelating reagent improves the reliability of IP isolation of EVs from biofluids. These results may reflect the fact that matrix proteins in specimens are associated with EV surface proteins (8), and these proteins inhibit interactions between antibodies and their targeted membrane proteins on EVs for IP assays.

Quantitative proteomic analysis revealed that IP using EDEG reduced the amount of calcium-dependent adhesion proteins including complement family proteins and immunoglobulin families (Supplementary Table.1). It has been reported that membrane adhesion proteins such as integrin, albumin, and cadherin family need metallic ions for interaction with proteins on the surface of the membrane (11). Since the EV marker protein CD9 is known to interact with integrin families (α6β1 and α5β1), fibronectin, and immunoglobulins (12), chelating reagent enables the dissociation of extracellular contaminant proteins and enhance the interaction between EVs surface CD9 and anti-CD9 antibody.

Furthermore, the quantitative proteomic analysis showed that the reduction of contaminant proteins increased the number of identified proteins, and EV marker proteins such as CD9 and CD81 were only identified in EDEG-based IP (Fig.3A). It indicates that extracellular matrix proteins abound in EVs fraction and they mask the proteins of interest when carrying out proteomics analyses. UC method is time-consuming and it includes substantial amount of contaminant proteins, while polymer method is rapid but additives or ingredients contained in EV isolation reagents sometimes adversely affect subsequent analyses, e.g., mass spectrometry. Our study showed that the EDEG chelating reagent is applicable to proteomic analysis as well as other omics analyses.

Taken together, our results show that chelating reagents are capable of efficiently recovering CD9 positive EVs from serum, plasma, and urine, which could contribute to clinical application of EVs and biomarker detection for diagnosis. EDEG combined with IP using an antibody against specific protein may even enable the isolation of certain EVs population, e.g., ones containing organ-specific membrane protein of clinical interest. It will also provide a reliable tool in clinical labs with automated systems.

## Acknowledgments

We acknowledge Ms. Haruka Shitara (PERSOL TEMPSTAFF CO., LTD.) for her technical support. We also thank Mr. Takuya Sakyu for the preparation of antibodies against CD9 and CD63.

## Figure legends

**Supplementary Table.1.**
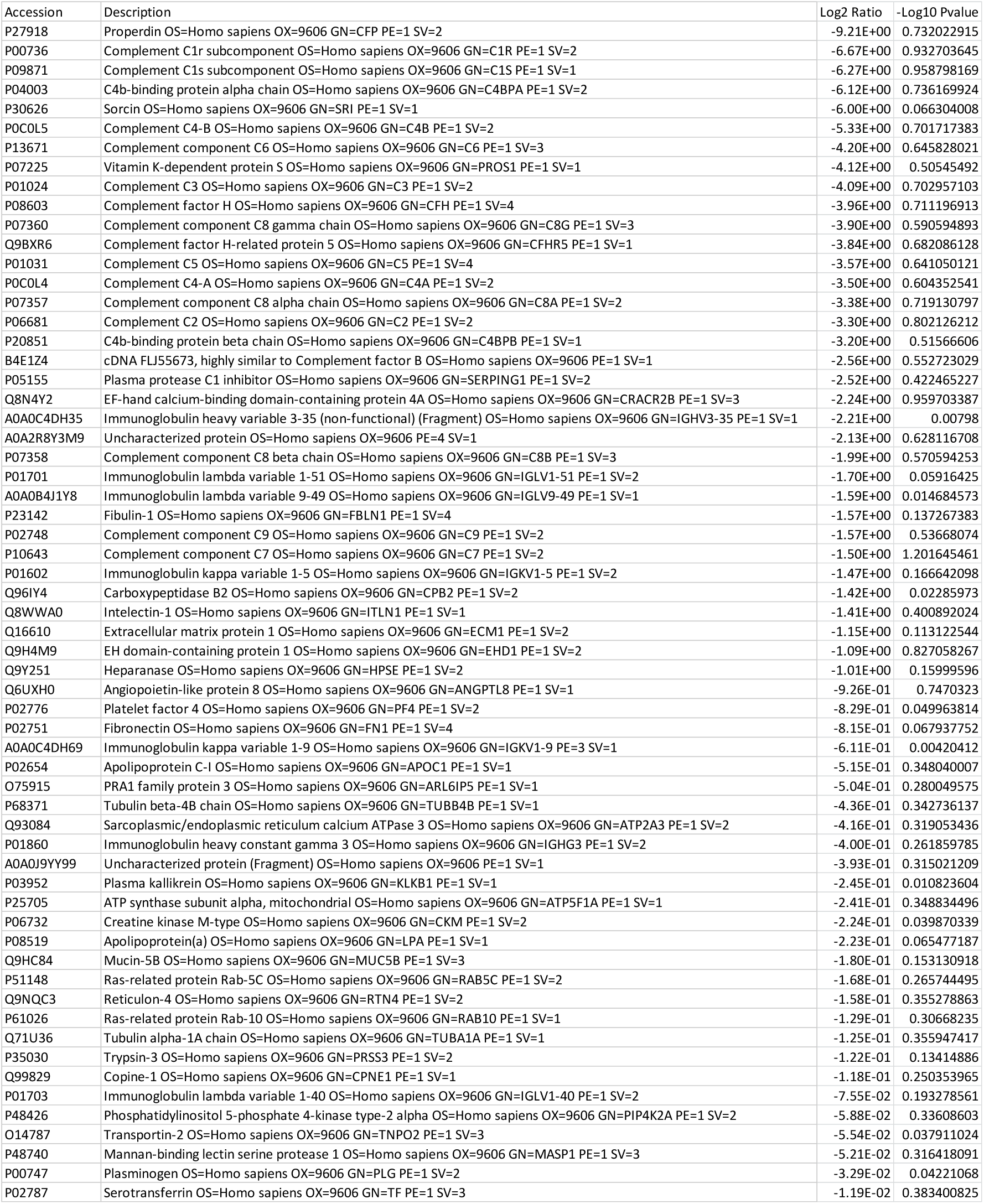
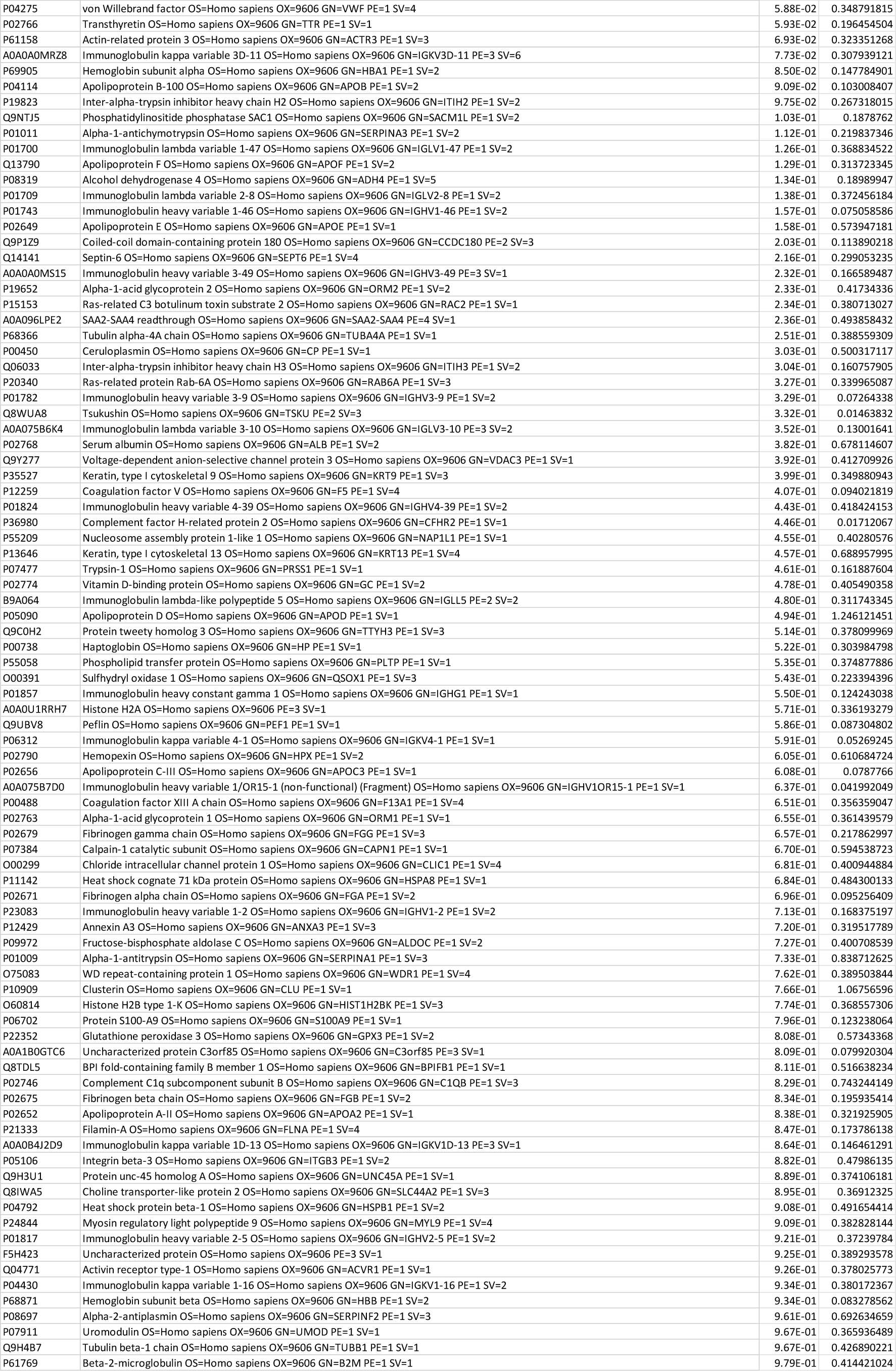
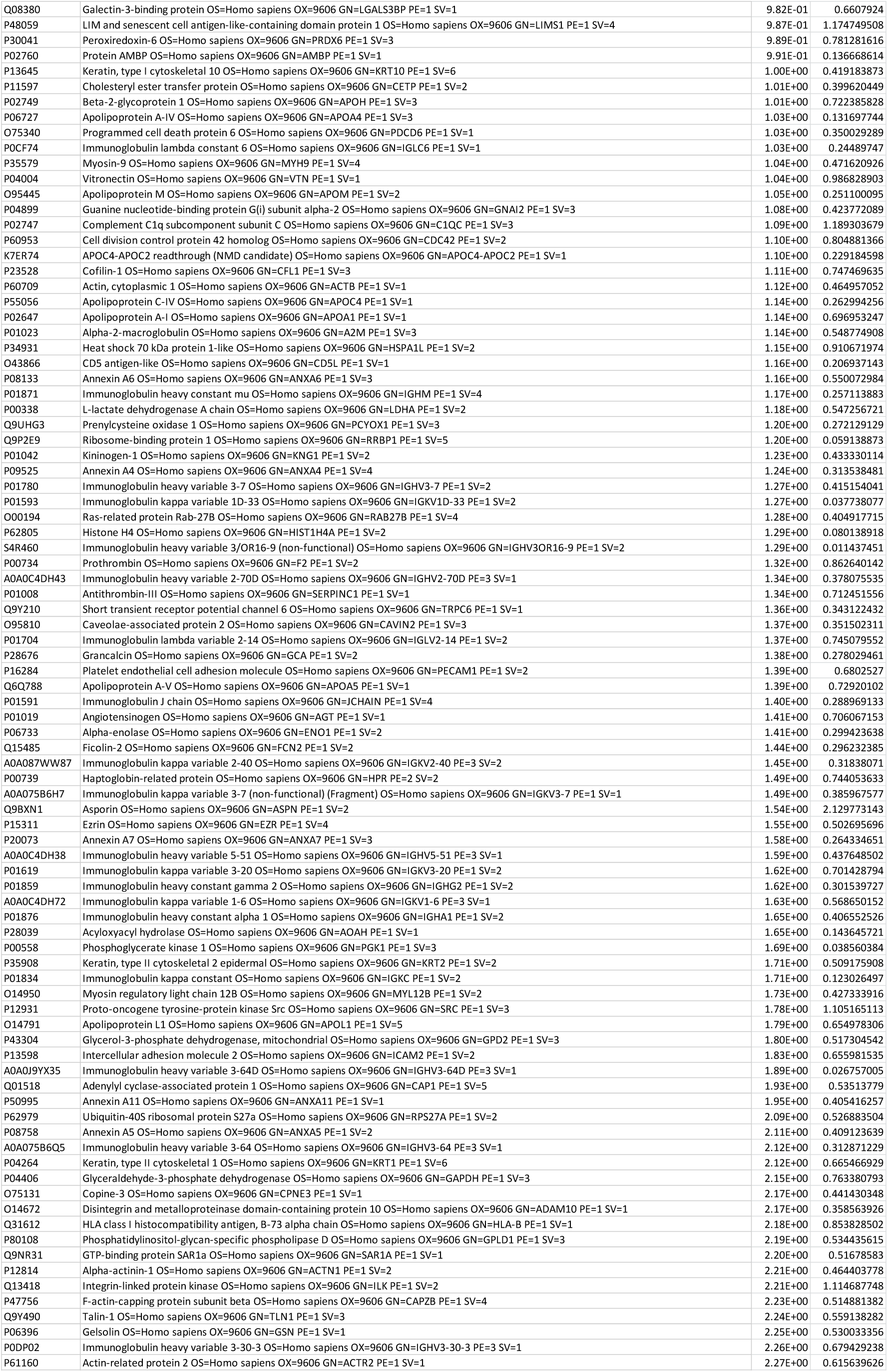
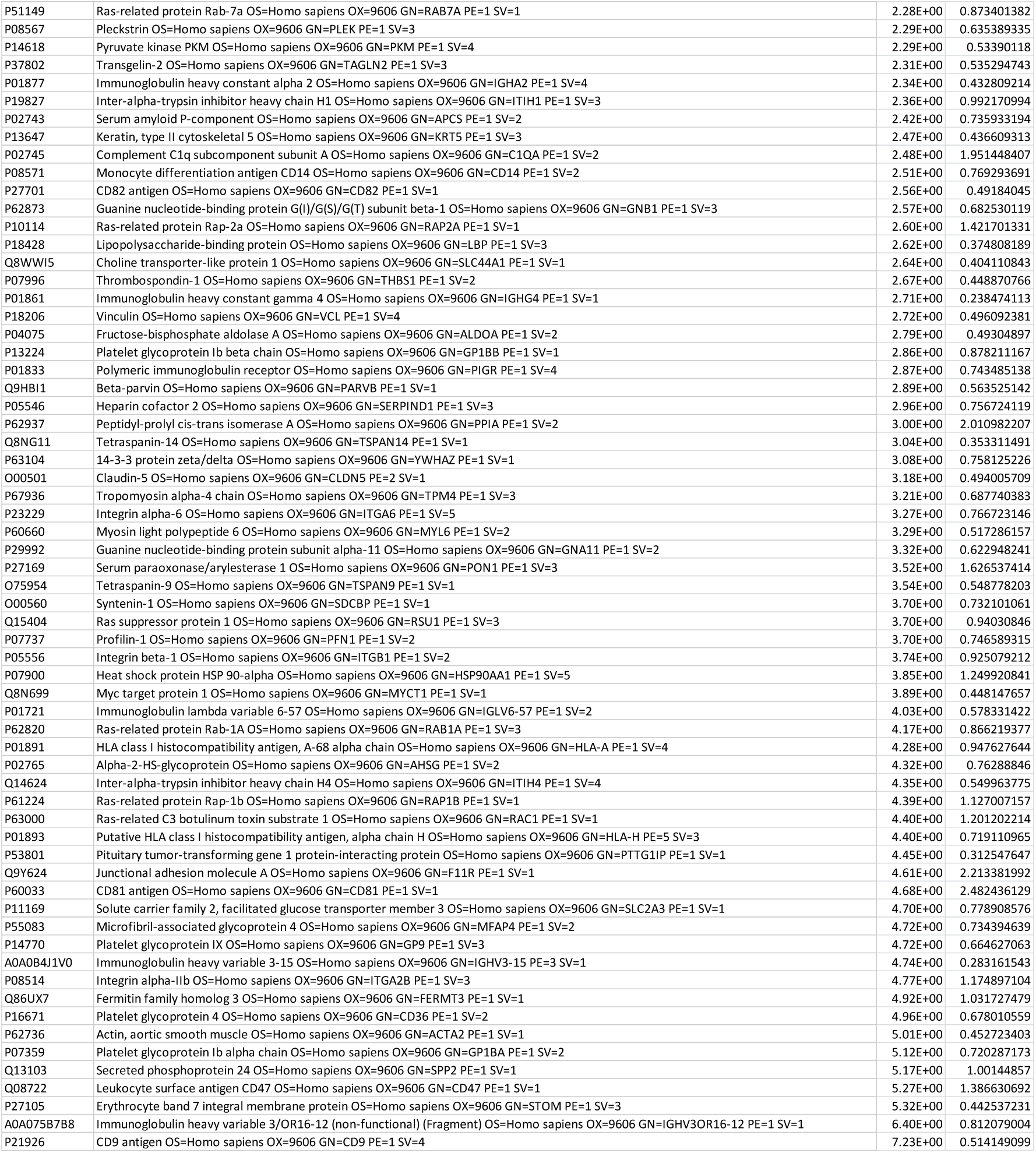
List of proteins identified in volcano plot analysis.

## Author contributions

Conceptualization: Ayako Kurimoto, Tatsutoshi Inuzuka.

Data curation: Ayako Kurimoto,

Investigation: Tatsutoshi Inuzuka, Yuki Kawasaki, Ayako Kurimoto, Toshiki Ueda.

Methodology: Tatsutoshi Inuzuka, Yuki Kawasaki, Ayako Kurimoto

Visualization: Ayako Kurimoto

Writing – original draft: Ayako Kurimoto

Writing – review & editing: Ayako Kurimoto, Tatsutoshi Inuzuka, Yuki Kawasaki, Fumi Asai, Koichiro Murashima, Kazuya Omi.

AK and TI conceived the project and designed the research. TI, YK, TU, and AK performed the experiments for isolation and characterization of EVs. AK wrote the paper. All authors read and approved the final manuscript.

